# IsarPipeline: Combining MMseqs2 and PSI-BLAST to Quickly Generate Extensive Protein Sequence Alignment Profiles

**DOI:** 10.1101/2022.03.23.485035

**Authors:** Issar Arab

## Abstract

Many of the machine learning (ML) models used in the field of bioinformatics and computational biology to predict either function or structure of proteins rely on the evolutionary information as summarized in multiple-sequence alignments (MSAs) or the resulting position-specific scoring matrices (PSSMs), as generated by PSI-BLAST. The current procedure used in protein structure and function prediction is computationally exhaustive and time-consuming. The main issue relies on the PSI-BLAST software being forced to load the current database of sequences (about 220 GB) in batches and search for similar sequence alignments to a query sequence. This leads to an average runtime of about 40-60 min for a medium-sized (450 Amino Acids) query protein. This average runtime is strictly dependent on the hardware used to run the software. The issue is becoming more problematic since the bio-sequence data pools are increasing in size exponentially over time, hence raising PSI-BLAST runtime as well. A prominent solution claims to speed up the current process by 100 folds. The MMseqs2 method, given enough memory, will load the whole database in memory and apply certain heuristics to retrieve the relevant set of aligned sequences. However, this solution cannot be used directly to generate the final output in the desired PSI-BLAST alignment and PSSM profile data format. In this research project, we analyzed the runtime performance of each tool separately. Furthermore, we built a pipeline that combines both MMseqs2 and PSI-BLAST to obtain a robust, optimized and very fast hybrid alignment tool, faster than PSI-BLAST by two orders of magnitude. It is implemented in C++ and is freely available under the MIT license at https://github.com/issararab/IsarPipeline. The output of our pipeline was evaluated on two previously built predictive models.

## I. Introduction

Multiple sequence alignment methods, or MSA-methods, are a set of algorithmic solutions for the alignment of evolutionarily related sequences. They can be applied to DNA, RNA, or protein sequences. Those algorithms are designed to take into account evolutionary events such as mutations, insertions, deletions, and rearrangements under certain conditions [1]. In their recent Nature publication, Van Noorden R. et al. [2] mentioned that in the field of biology, the most widely adopted modeling approaches use multiple sequence alignment methods. This statement is supported by Altschul et al. [3] in their published manuscript of the tool “Gapped BLAST and PSI-BLAST,” that is ranked the 11th most cited scientific paper of all times [2]. Indeed, the development of this vital modeling tool, MSA, necessitated the addressing of both complex computational and biological problems. MSA has always been recognized as an NP-complete problem. That is the reason behind the numerous alternative algorithms built, more than 100 variants, aiming at providing an accurate MSA over the last four decades [4].

In contemporary computational biology and, more specifically, in the area of proteins and the prediction of their properties, sequence alignment forms the de-facto standard input to nearly all ML methods [5, 6]. However, this methodology is computationally exhaustive and time-consuming.

Currently, MMseqs2 [7, 8] appears to be the only MSA-based alternative solution out there coping with these enormous datasets, but at the expense of substantial hardware requirements. We introduce in this paper an optimized and fast hybrid alignment solution combining these two state-of-the-art methods to quickly generate extensive protein sequence alignments and corresponding PSSM profiles. In this manuscript, we explain the pipeline architecture components, give a runtime comparison between both tools (MMseqs2 vs PSI-BLAST), and finally, present the evaluation results of the IsarPipeline output with regard to two ML models (TMSEG [16] and REPROF [17]) from Predict-Protein [15].

All results presented in the following sections were conducted on a server granted by the University of Luxembourg. The machine’s main characteristics are summarized as follows: a memory size of 196Gib with an Intel Xeon E312xx CPU of 4 cores supporting the SSE4.1 instruction set.

## II. Background

The three main elements of IsarPipeline are: MMseqs2, PSI-BLAST, and the generated PSSM profile.

### A. MMseqs2

MMseqs2 [7, 8] stands for Many-against-Many Sequence Searching. It is an extremely optimized scalable software used to search and cluster very large protein sequence datasets. It is an open-source cross-platform tool developed in C++ using the full power of modern CPU architectures, especially the SIMD optimized instruction set processing unit. The tool implements a SIMD accelerated Smith-Waterman-alignment algorithm [12] to compute the alignment of protein sequences.

### B. PSI-BLAST

PSI-BLAST [9] stands for the Position-Specific Iterative Basic Local Alignment Search Tool. It is a software that runs a multiple sequence alignment algorithm powered by the dynamic programming optimization paradigm to search a given database of protein sequences [6]. Given a query sequence, the algorithm retrieves homologous sequences that pass a predefined threshold. This threshold-based computational similarity method, which uses protein-protein BLAST [6, 9, 10] to identify regions of local alignment, is used to construct a PSSM from the retrieved family of sequences. The matrix is used as input for subsequent iterations of the software to further search the database and retrieve more hits, which are then used to update and improve the PSSM itself.

### C. PSSM

A PSSM stands for the Position-Specific Scoring Matrix, which is also referred to as a profile matrix. The PSSM profiles contain statistical representations of residues in a given sequence of a protein with respect to all its relevant aligned protein sequences (i.e. homologs) in the database. A profile captures the conservation pattern in an alignment and stores it as a matrix of scores for each position in the alignment [9], where high scores are assigned to highly conserved positions and lower scores to low conserved ones. Hence, the matrix representation includes what we call the protein evolutionary information. PSSMs prove themselves to be essential for the field of bioinformatics and computational biology. However, the calculation of those PSSMs is not affordable since it requires significant computation power and runtime. This computational cost is well noticed during the building process of a new ML method or any statistical analysis which both require thousands of samples to be built and evaluated to achieve good results. Those samples will need a few weeks of processing before you can start your study, which makes it highly inconvenient.

## III. Dataset

Runtime measurements in this paper refer to search alignments of amino acid sequences (excluding the time needed for indexing ~1h20min) against a sample of UniProt [11] Reference Cluster with 90% sequence identity (uniref90 2019_02). For faster execution, we need an index table of the database that can fit into the RAM. Since the whole uniref90 will generate an index table of roughly the size of 250Gib, we run our tests on a random sample (~ 68 million proteins) from uniref90. The sample generates 181Gib index table, which can fit in our test machine memory.

## IV. IsarPipeline Architecture

IsarPipeline combines MMseqs2 and PSI-BLAST to quickly generate profiles for a set of query sequences using the heuristics of the first method, which aligns batch query sequences in parallel, and the ability of the second tool to generate corresponding PSSM profiles. To achieve this goal, a pre-processing of the database to be searched is crucial before running IsarPipeline (Fig. 1). The pre-processing step generates an index table, which can be of a magnitude of 100s of gigabytes for big datasets. This step is essential to make the alignment faster afterwards. Creating an index table is not mandatory to search a database via IsarPipeline. The search command will create the index table right before starting the pipeline execution. However, it is highly recommended to generate the index table and store it beforehand. This way, the search command will not construct one for each search, especially if one is starting multiple runs of the pipeline on the same database.

**Fig. 1.**
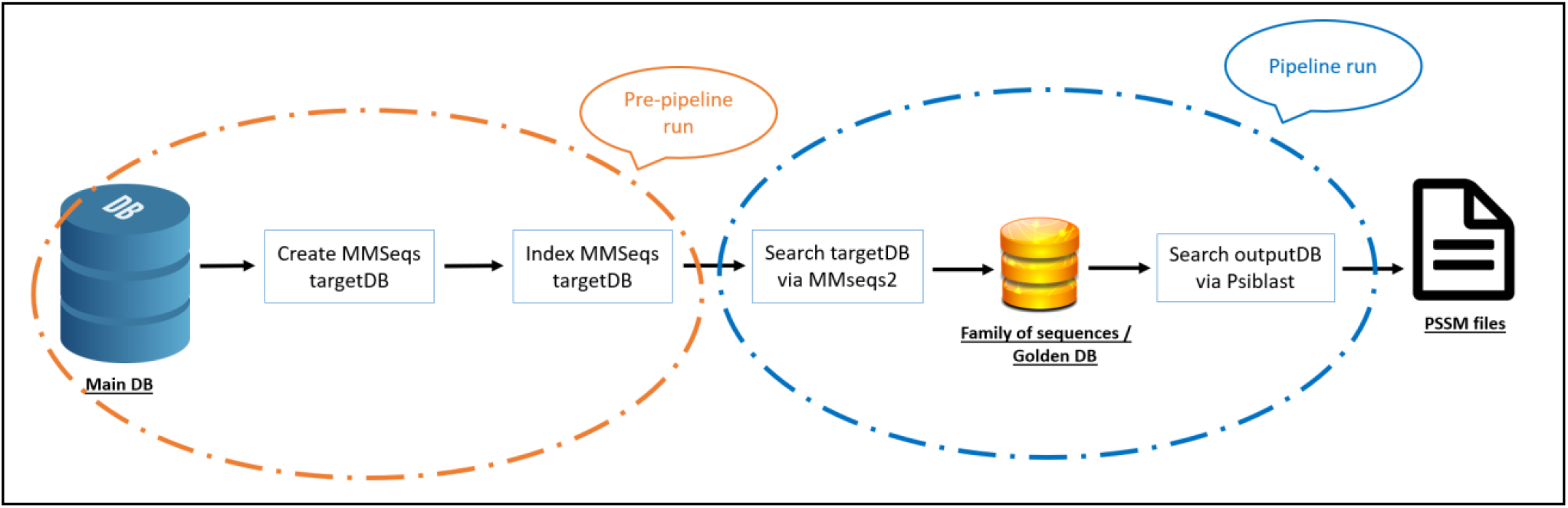
IsarPipeline high-level architecture. The high-level architecture consists of two main parts: a pre-processing step, orange section, and a pipeline execution step, blue section. The pre-pipeline procedure compiles an MMseqs database, named targetDB, from the FASTA file of the main database to be searched. Then, from the targetDB we construct a big index table that later will be used by the pipeline for fast query alignment. This pre-pipeline step is crucial for a faster and reusable index table in future runs.

The output of the pre-processing step (Fig. 1 - orange) is an index table of the main target database to be searched. It represents the input to IsarPipeline (Fig. 1 - blue), which runs a set of consecutive modules in order to finally generate a PSSM profile. The first one to kick-off is the “pre-filter module”. This module constructs the main component of MMseqs2 power. The pre-filtering module computes an ungapped alignment score for all consecutive *k-mer* matches between all query sequences and all database sequences and returns the highest score per sequence [4, 5, and 8]. The prefilter *k-mer* match stage is key to the high speed and sensitivity. It detects consecutive short words (i.e. *k-mer*) matching on the same diagonal. The diagonal of a *k-mer* match is the difference between the positions of the two similar *k-mer* in the query and in the target sequence [4, 5, and 8]. Once the pre-filtering module is complete, it generates the best matches for each query sequence. Those matches do not go beyond the number *N*, the max number of sequences to be retrieved by the module. This number can be provided as an argument to the pipeline and is set in our experiments to 1000 aligned sequences per query sequence.

The hits are then passed to the “alignment module”. This module implements a SIMD accelerated Smith-Waterman-alignment algorithm [12] of all sequences that pass a cut-off for the prefiltering score in the first module. It processes each sequence pair from the prefiltering results and aligns them in parallel, calculating one alignment per core at a single point of time. This step is implemented using a vectorized programming algorithm that makes use of the SIMD instruction set. This is where the SSE4.1 instruction set requirement is being used. In the end, this module calculates alignment statistics such as sequence identity, alignment coverage and e-value of the alignment [13], which are then passed to the “convertalis module” and the “parsing module” to transform the output into FASTA format, using optimized algorithms detailed in [14]. We call this output the golden DB with a maximum of 1k sequences, which are then fed to PSI-BLAST modules in order to generate the final PSSM profile (Refer to section 1 in the SOM for detailed component description with a schematic).

## V. Super-Fast Sequence Alignment

Analysis of each module runtime in IsarPipeline search revealed that loading (I/O) the index table represents ~90% of the whole runtime reported, which is mainly constrained during the pre-filtering and the Convertalis module execution. Therefore, a way to load the whole DB into RAM beforehand, then use it directly during each search, as well as the conversion of the search results, will decrease our runtime tremendously.

One hack is to load the whole database into RAM and lock the pages there. Later in the search, we can map the sequences from the locked DB instead of loading it again from disk. We used a well-maintained software called “vmtouch”, a tool for learning about and controlling the file system cache of Unix and Unix-like systems [18]. The tool loads the index table into RAM and lock the pages there. You can run it via the following command:

> ***sudo /usr/local/bin/vmtouch -l -d -t targetDB***.***idx***

The command has two main parameters: “-l”, to lock the pages, and “-d”, to run it as a daemon. Also, the command must have root privileges. The reason behind that is the “max lock memory” constraint enforced by the operating system. It is usual 64kb. Otherwise, your database will be loaded into RAM but not locked there. This can be resolved by either including the full PATH into *sudo* environment variables and running the process with root privileges or just running the command with *sudo* but using the full path to the executable.

With this hack and providing “--db-load-mode 2” as a parameter value, we were able to generate the PSSM file for an average length protein sequence in 53 secs using IsarPipeline.

## VI. Evaluation and Results

### A. MMseqs2 VS PSI-BLAST: Runtime Benchmarking

Comparing the runtime execution of each of the methods separately using a single query protein shows that the amino acid sequence length of the protein affects PSI-BLAST (Fig. S2 in SOM) runtime while it does not for MMseqs2. We observe that PSI-BLAST runtime increases with the increasing number of residues in a query sequence while MMseqs2 runtime remains constant. However, the experiments show the opposite for batch processing. We notice that the increasing number of sequences included in one query heavily affects PSI-BLAST runtime (Fig. S3&2). On the other hand, MMseqs2 is barely affected (Fig. 2-b). MMseqs2 runtime ranges between 22min and 32min with an average runtime of ~30min. PSI-BLAST runtime increases linearly, and this is mainly due to the sequential processing of the query sequences.

**Fig. 2.**
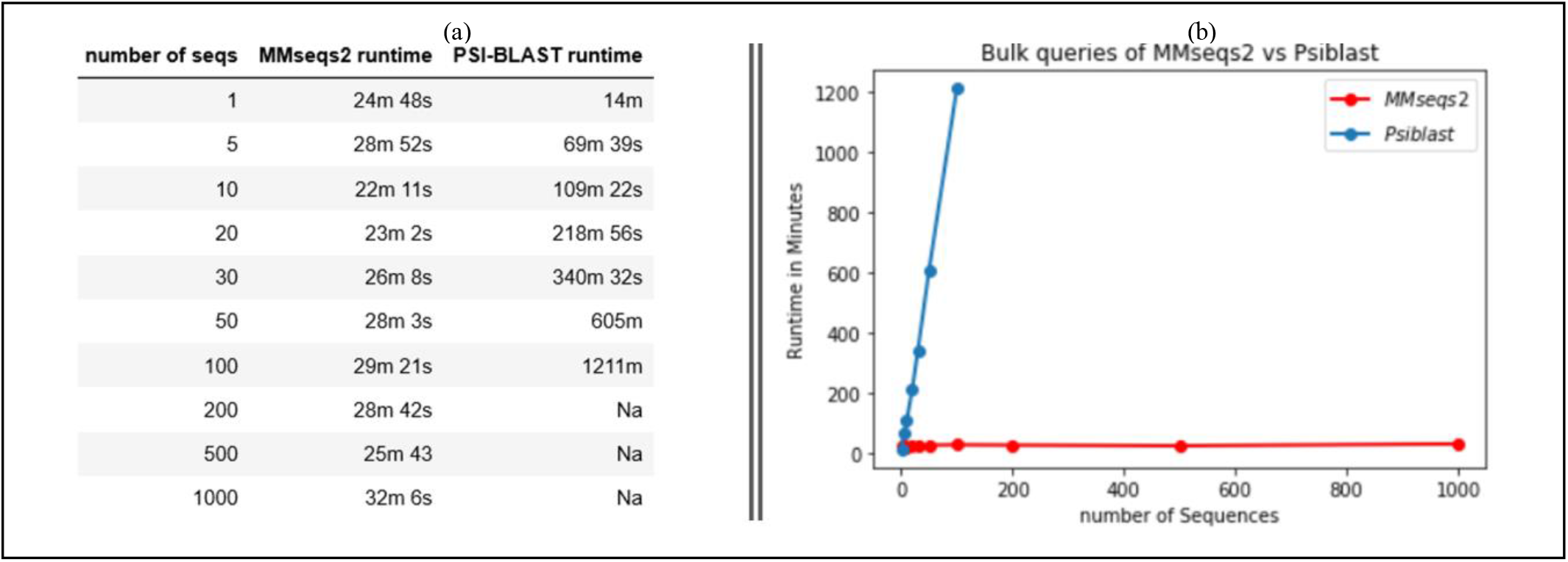
(a) table representing the effect of random bulk sequences as a single query on the runtime of both MMseqs2 vs PSI-BLAST. (b) is a visual representation of table (a) comparing the effect of the number of sequences on the runtime of each alignment software. PSI-BLAST scales linearly, blue, while MMseqs2 remains relatively constant, red

PSI-BLAST is faster when it comes to a single query alignment while MMseqs2 scales much better in terms of batch sequence processing. This feature was exploited to build a faster batch sequence alignment method.

### B. Pipeline Evaluation

One way to evaluate the output results of our IsarPipeline is to assess how the predictions of our existing ML models using our tool output are changing and whether the pipeline has a negative effect on the predictions. We tested the pipeline on two ML methods: TMSEG [16] and REPROF [17].

#### 1) TMSEG

TMSEG is a tool that predicts the transmembrane segments in a protein. For this method, 265 unique proteins were selected to be evaluated using both PSI-BLAST and IsarPipeline. The first metric dimension to evaluate is the runtime needed to get the PSSM files. PSI-BLAST takes 48.5h (Tab. 1) to generate the PSSMs while IsarPipeline needs only ½ h, a drastic decrease in runtime by ~100 folds.

**Table I.** Runtime comparison of running 265 query batch against a random sample of 68 million proteins from uniref90. PSI-BLAST needs 48h 28m, roughly 100 times more than IsarPipeline 31m, to generate PSSM profiles.

| number of seqs | IsarPipeline runtime | PSI-BLAST runtime |
| --- | --- | --- |
| 256 | 31m 13s | 48h 28m 11s |

The second metric dimension to evaluate is the characterwise similarity between both outputs. Fig. 3-b shows that the number of instances with a characterwise similarity percentage to the ground truth greater or equal to 0.95 resulting from PSI-BLAST PSSMs are lower than the ones resulting from IsarPipeline, as shown in Fig. 3-a. However, PSI-BLAST results with 0.9 similarity are much higher than IsarPipeline results. From a statistical point of view comparing the overall similarity means, IsarPipeline results have a similarity mean of 0.830 with a standard deviation (STD) of 0.125, whereas PSI-BLAST results have a similarity mean of 0.832 and a STD of 0.119. Therefore, both tools have roughly similar performance. The distribution of instances among all bars compensates for the differences in some regions. In terms of the predicted string using PSSMs from both tools, Fig. 4 shows that most of the predicted pairs are the same with a similarity mean of 0.945 and a STD of 0.061.

**Fig. 3.**
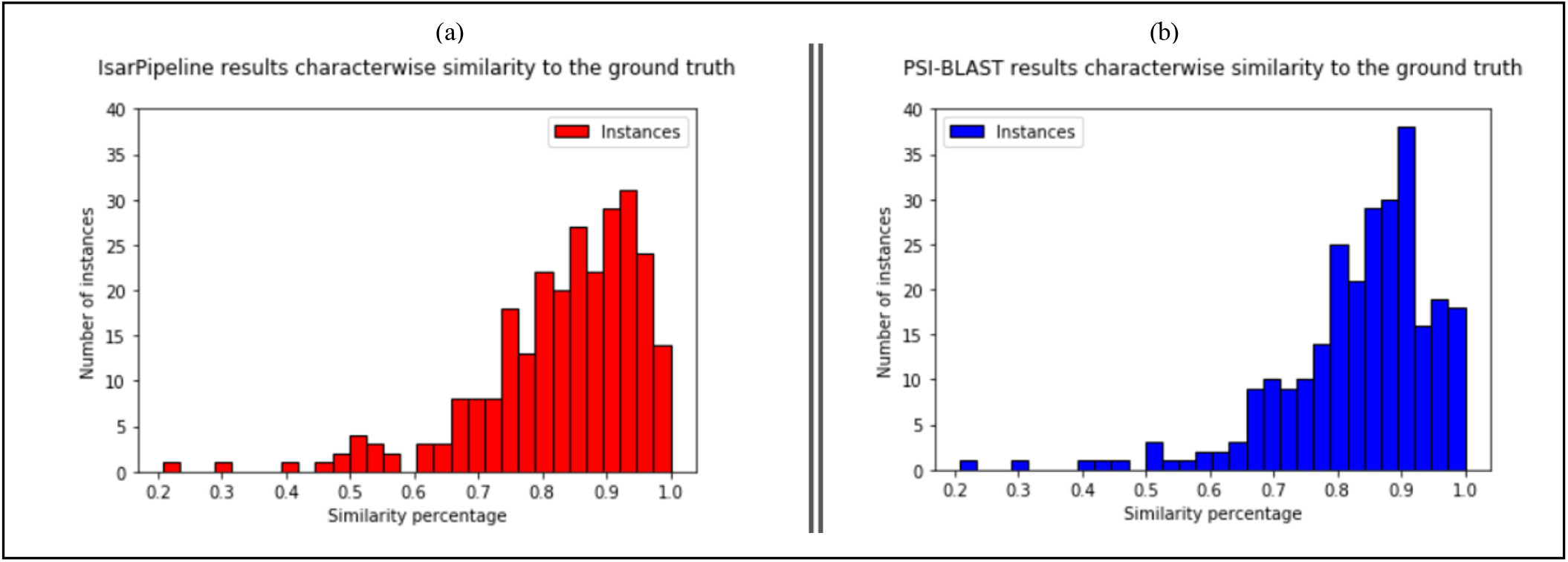
Characterwise TMSEG prediction similarity to the ground truth. Histogram (a) represents the percentage distribution of how similar the residue predictions of a sequence are to the ground truth labels using IsarPipeline. Histogram (b) represents the percentage distribution of how similar the residue predictions of a sequence are to the ground truth labels using PSI-BLAST.

**Fig. 4.**
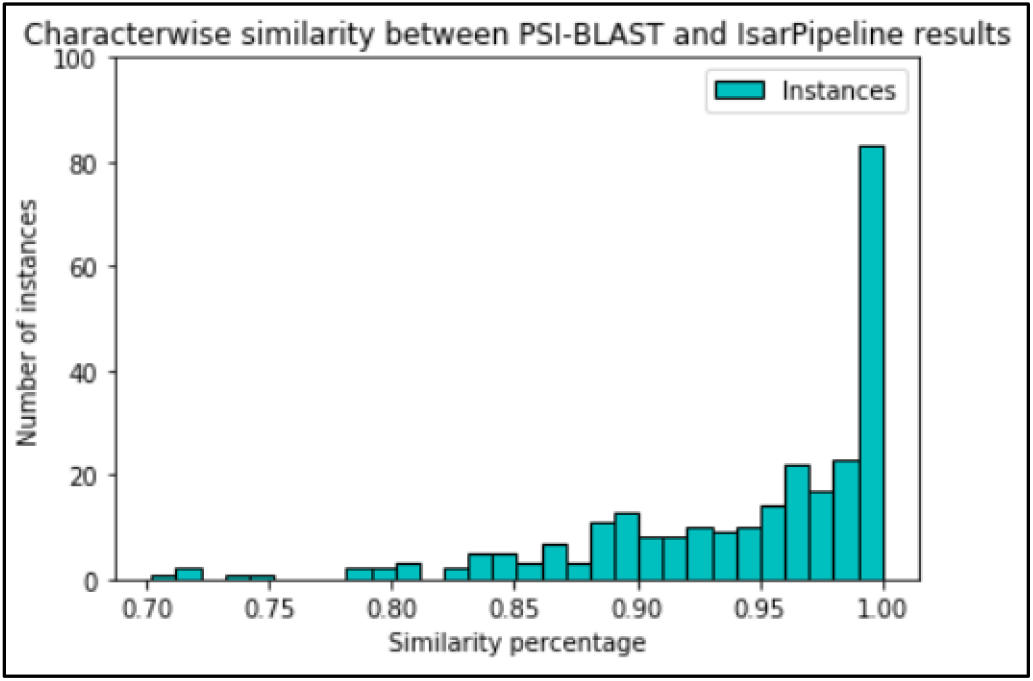
Characterwise percentage similarity between PSI-BLAST and IsarPipeline PSSM based predicted strings of the same amino acid sequence

To further analyze the output predictions generated based on IsarPipeline, we conducted a segment prediction analysis on the same test set and an extra analysis using the full dataset (8817 proteins). The results showed a similar trend, with a runtime execution of IsarPiepline 540 times less than PSI-BLAST (Tab. 2). For further details refer to section 3 of the SOM.

**Table II.** Runtime comparison of running 250 query batch against a random sample of 68 million proteins from uniref90. PSI-BLAST needs 50h 33m, roughly 100 times more than IsarPipeline 32m, to generate PSSM profiles.

| number of seqs | IsarPipeline runtime | PSI-BLAST runtime |
| --- | --- | --- |
| 250 | 32m 33s | 50h 33m 57s |

#### 2) REPROF

REPROF is a tool that predicts a Q3 protein secondary structure. For this tool, 250 unique proteins were selected to be evaluated using both PSI-BLAST and IsarPipeline. The first metric to evaluate for this ML method is the runtime needed to get the PSSM files. PSI-Blast takes 50.5h (Tab. 2) to get the PSSM files while IsarPipeline needs only ½h, a significant decrease in runtime by ~100 folds.

Similarly to TMSEG, a characterwise similarity analysis was conducted. The number of instances with a characterwise similarity percentage to the ground truth at 0.8 resulting from PSI-BLAST PSSMs (Fig. 5-b) are lower in magnitude than the ones resulting from IsarPipeline (Fig. 5-a). However, in general, both histograms look comparable with some bars compensating for other differences. This observation is supported by the statistics computed to compare the overall similarity means. IsarPipeline has a similarity mean of 0.781 with a STD of 0.092, whereas PSI-BLAST has a similarity mean of 0.779 with a STD of 0.092. Therefore, both tools have roughly similar performance. In terms of the predicted string using PSSMs from both tools, Fig. 6 shows that most of the predicted pairs are the same with a similarity mean of 0.0005 and a STD of 0.281.

**Fig. 5.**
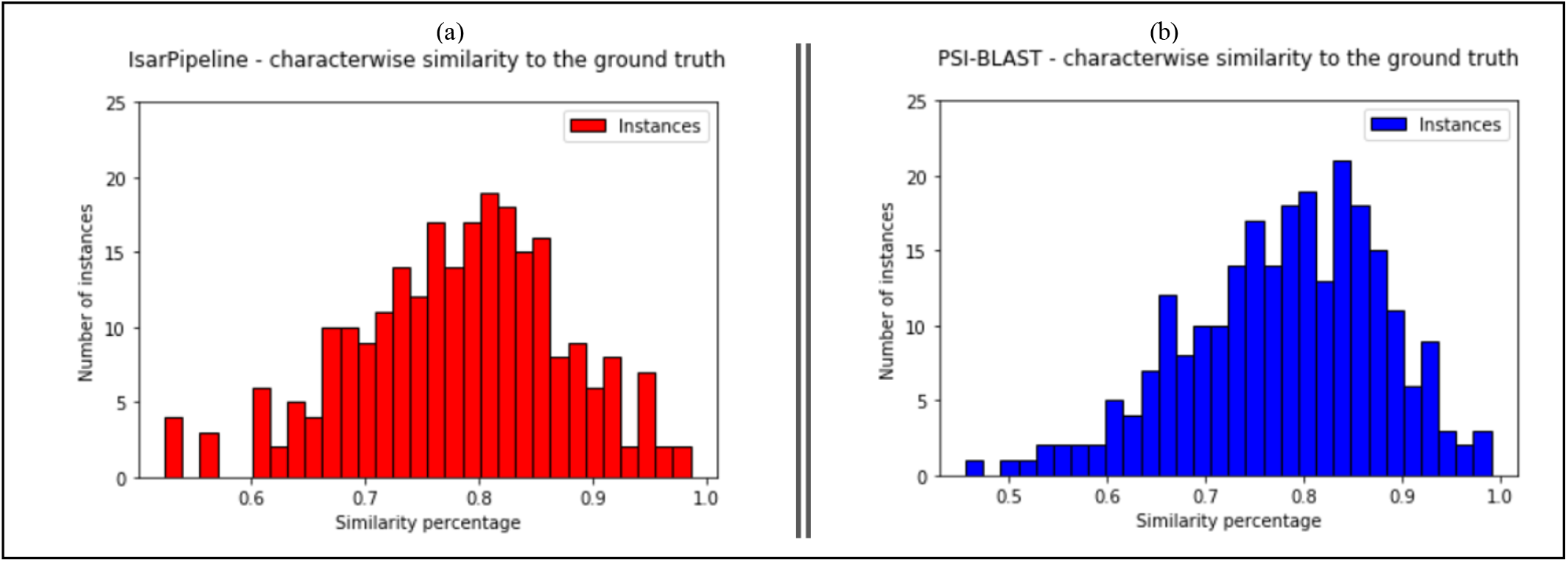
Characterwise REPROF prediction similarity to the ground truth. Histogram (a) represents the percentage distribution of how similar the residue predictions in a sequence are to the ground truth labels using IsarPipeline. Histogram (b) represents the percentage distribution of how similar the residue predictions in a sequence are to the ground truth labels using PSI-BLAST results.

**Fig. 6.**
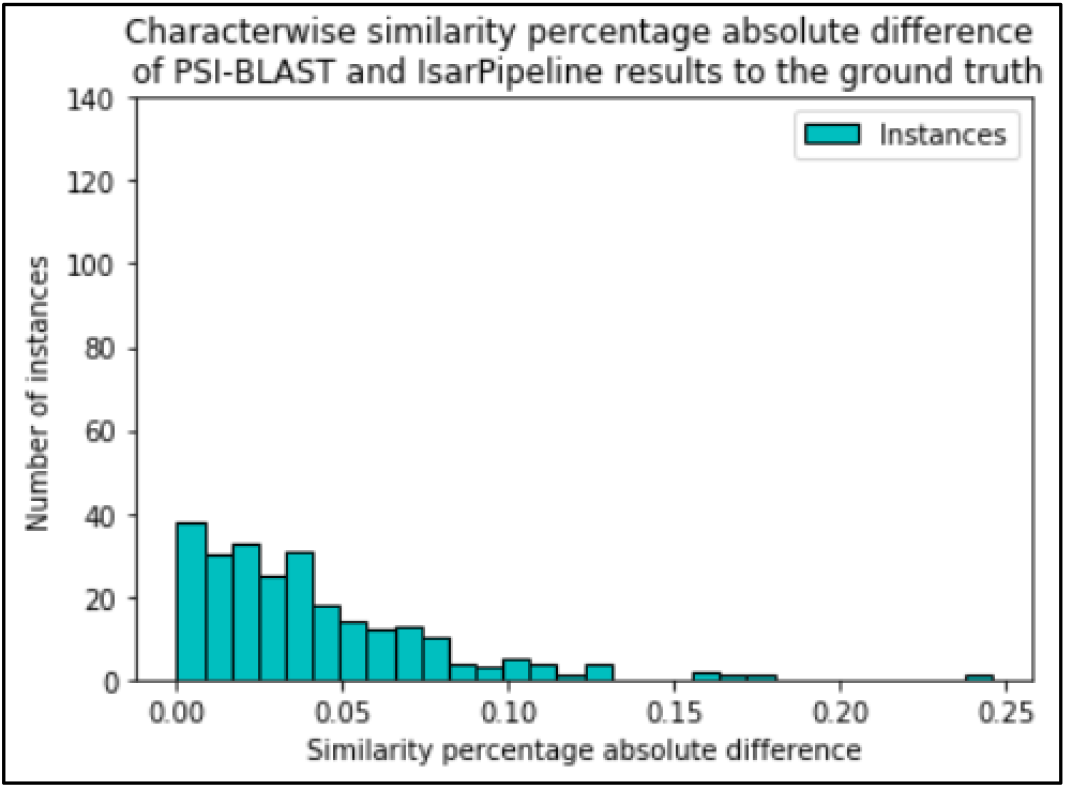
The histogram depicts the absolute difference of the characterwise percentage similarity between PSI-BLAST results and IsarPipeline results to the ground truth labels

A full dataset (4692 proteins) analysis was also performed and showed a similar trend, with a runtime improvement of 414 times. For further details refer to the SOM, section 4.

## VII. Conclusion

Results of this research work reveal that MMseqs2 runs slower when it comes to a single query search compared to PSI-BLAST. However, it demonstrates being powerful when it comes to batch processing. We have exploited this property to build a pipeline, IsarPipeline, that combines both MMseqs2 and PSI-BLAST to quickly generate extensive sequence alignments and construct the PSSM profiles. These profile matrices are used in most of the ML predictive models out there. When searching for large query batches of protein sequences, IsarPipeline performance proves to be faster by more than 2 orders of magnitude compared to PSI-BLAST. We have evaluated the output results of the pipeline and compared them to the ones of PSI-PLAST. We have seen that IsarPipeline performance is at least comparable to and often slightly better than the PSI-BLAST results. Later, we have introduced a hack to make a single query alignment almost instant. Instead of loading the database for each search, using “vmtouch”, one can load the index table into the RAM and lock its pages that will then be used directly through mapping of sequences for single query searches. Eventually, we achieved a runtime speed of 53 secs, for an average length sequence (~450 AA), with IsarPipeline. The alignment generates a PSSM file incorporating almost the same statistical information as the matrix generated by PSI-BLAST in tens of minutes.

## Acknowledgment

The author would like to thank Michael Bernhofer, Martin Steinegger, and Burkhard Rost for their insights and feedback on this project. The author would like to also thank the University of Luxembourg for the dedicated computing resources granted for this project.

