## supplementary online material for "IsarPipeline: Combining MMseqs2 and PSI-BLAST to Quickly Generate Extensive Protein Sequence Alignment Profiles"

#### 1- IsarPipeline Modules

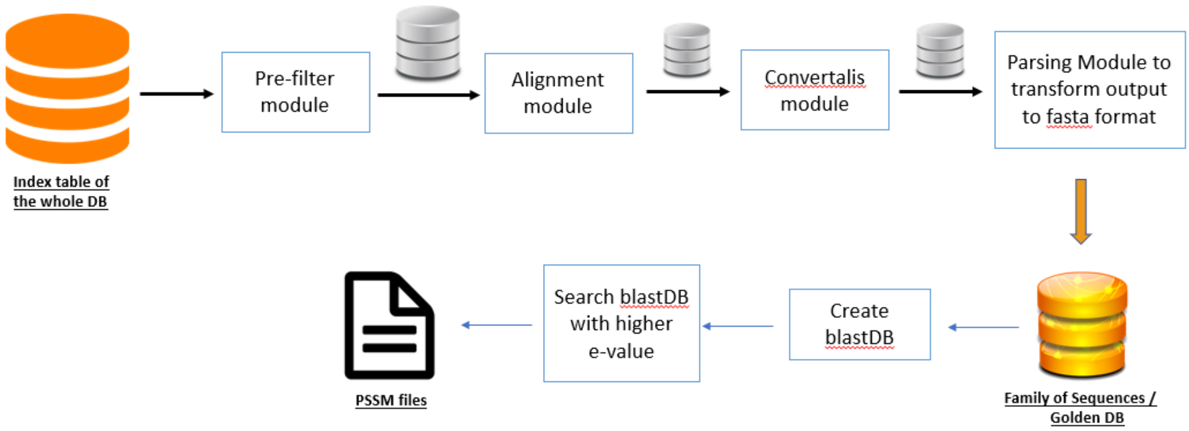


**Figure S1:** IsarPipeline Low-Level Architecture. The Pipeline requires an index table of the database to be searched as input then runs the pre-filtering (Pre-filter module) with a maximum number of sequences to be retrieved (a pipeline parameter that can be tweaked - set to 1000 in our experiments). The result is then aligned (Alignment module) to produce a resultdb that will then be converted (Convertalis module) to isar.tuple, a big file containing a tab-separated list of 3 columns: query header, target header, and target sequence. The file is then parsed (Parsing module) to create a query file and its corresponding db file, in FASTA format. The query and db files will then be converted to BLAST output (local alignment result, PSSM, or ASCII PSSM – pipeline parameters to be provided) via PSI-BLAST search on the parsed db with a high e-value (set to 10).

IsarPipeline takes an index table as input, then runs the pre-filter and alignment modules to allocate the best hits to a query sequence and compute a set of statistics. Output is then fed to the Convertalis module, which takes the alignment output and converts the result database into a BLAST tab formatted file. The structure of the file is a tab-separated list with 3 columns. The first column includes the header of a query sequence. The second column consists of the header of the target sequence. The third column has the amino acid sequence of the target protein.

The third step results in a very large file containing all the results. This file can reach 1 million lines for a query batch of 1000 query sequences. The main purpose of the parsing module is to split the output file of the Convertalis module into a list of query files and their corresponding targetDB files. For instance, if the query consists of one protein sequence, the output of this module should be two separate files, a query file and a DB/target file. Then if the query consists of 1000 sequences, the output will be 1000 query files each containing a sequence and 1000 corresponding DB/target files. The processing becomes more critical with a higher number of query sets. Hence, an optimal solution was to load the whole output file into memory using a hack explained in the book Optimized C++, Proven Techniques for Heightened Performance [14]. Basically, the idea is to take bigger bites using bigger input buffer. C++ streams contain a class derived from std::streambuf that improves performance in the case of file reading by reading data from the underlying operating system in bigger chunks [14]. Once loaded, the file content is tokenized using optimized C++ algorithms from the boost library maintained by IBM.

Once we have the family of sequences (named Golden DB in Fig. S1), it is time to run PSI-BLAST to generate the profiles and PSSM files. In order to scale up this module efficiently, a parallelized implementation is deemed necessary. Hence, the pipeline detects automatically the number of cores available in the machine and dispatches the jobs in a stratified manner, in such away each core will process the same number of queries at the same time. At this stage, PSI-BLAST is run with a higher e-value to capture all the results produced by previous steps.

### 2- Runtime Benchmarking

(a)

| seqlen | MMseqs2 runtime | PSI-BLAST runtime |
|---|---|---|
| 98 | 30m 24s | 12m 51s |
| 212 | 24m 35s | 14m 8s |
| 543 | 24m 20s | 17m 15s |
| 795 | 24m 48s | 14m 20s |
| 1040 | 24m 3s | 15m 35s |
| 1553 | 24m 8s | 22m 10s |

(b)

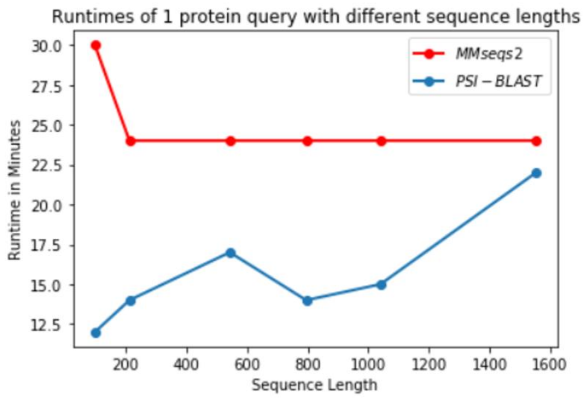


**Figure S2.** (a) table representing runtime needed to run one query with sequence length (seqlen) against sample uniref90 with each software alone (MMseqs2 or PSI-BLAST) (b) graph depicting the effect of sequence length (x-axis) on the runtime (y-axis) using each alignment method. Sequence length affects PSI-BLAST runtime while it does not for MMseqs2.

(a)

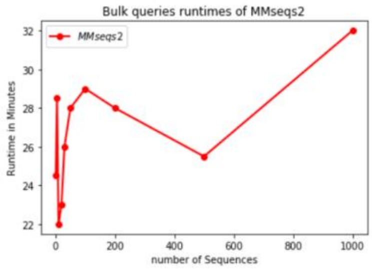


(b)

| number of seqs | MMseqs2 runtime | PSI-BLAST runtime |
|---|---|---|
| 1 | 24m 48s | 14m |
| 5 | 28m 52s | 69m 39s |
| 10 | 22m 11s | 109m 22s |
| 20 | 23m 2s | 218m 56s |
| 30 | 26m 8s | 340m 32s |
| 50 | 28m 3s | 605m |
| 100 | 29m 21s | 1211m |
| 200 | 28m 42s | Na |
| 500 | 25m 43 | Na |
| 1000 | 32m 6s | Na |

(c)

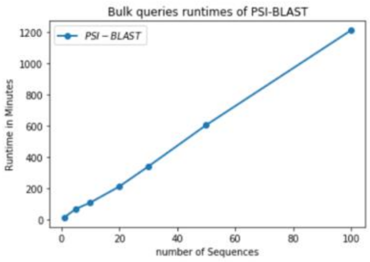


**Figure S3.** (b) table representing the effect of random bulk sequences (number of seqs) as a single query on the runtime of MMseqs2 in (a) and PSI-BLAST in (c).

### 3- IsarPipeline Evaluation on TMSEG

#### 3.1- Segment Prediction Analysis

TMSEG segment-based predictions represent the predicted regions (a segment of consecutive residues) of a query sequence where those segments match the transmembrane helix regions of the ground truth.

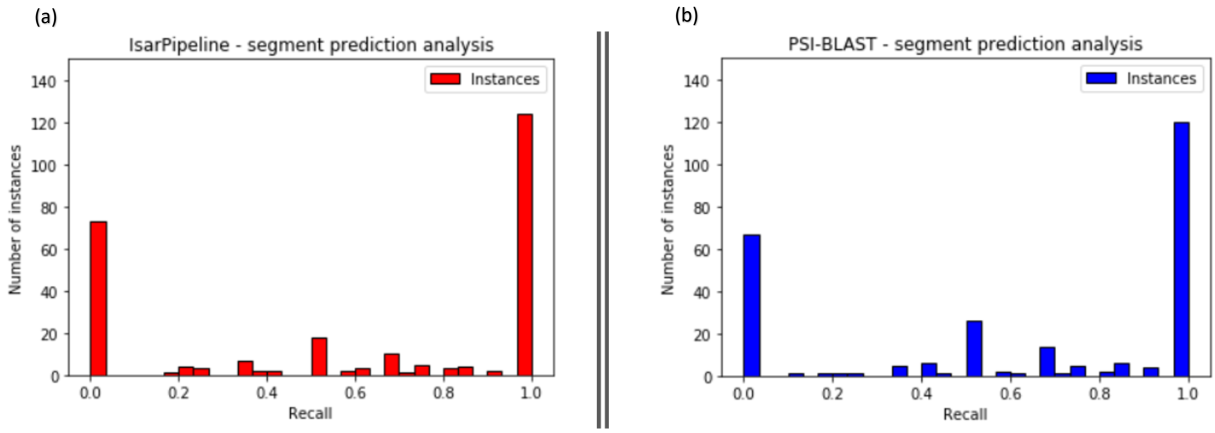


**Figure S4**. Segment-based TMSEG prediction analysis to the ground truth. Histogram (a) represents the recall distribution, which is the percentage distribution of how many segments in a sequence ground truth are correctly predicted using IsarPipeline results. Histogram (b) represents the recall distribution, which is the percentage distribution of how many segments in a sequence ground truth are correctly predicted using PSI-BLAST results.

The histogram distribution of the recall metric (Fig. S4) shows that we have a similar number of correctly predicted segments from both tools. Also, half of the instances (Fig. S4), segments in a particular instance/sequence, are 100% correctly predicted. Furthermore, from a statistical point of view and comparing the overall similarity means, IsarPipeline has a recall mean of **0.607** with a STD of **0.430.** Whereas PSI-BLAST has a recall mean of **0.621** with a STD of **0.416**. We observe that both tools have again roughly a similar performance, but are the correctly predicted results the same in both PSI-BLAST and IsarPipeline? To do that we compute the absolute difference mean between IsarPipeline and PSI-BLAST prediction output pairs.

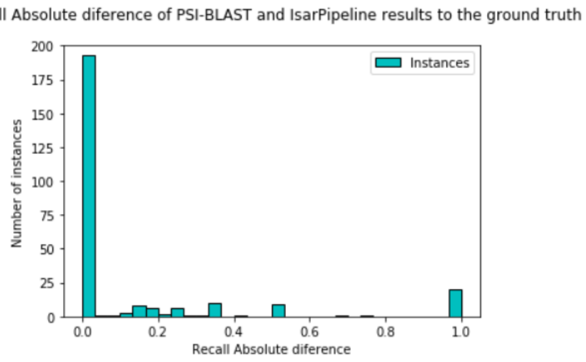


**Figure S5**. Segment recall absolute difference analysis using TMSEG between PSI-BLAST and IsarPipeline PSSM based predicted strings of the same amino acid sequence

The analysis demonstrates (Fig. S5) that more than 80% of the recall of predicted pairs are the same with an absolute difference mean of **0.0005** and a STD of **0.281**.

#### 3.2 - Full Dataset Analysis

To make sure that the sample we tested on is representative, we check whether the statistics computed previously still generalize for the whole TMSEG dataset, without redundancy reduction, of about 8817 protein sequences. Since running PSI-BLAST on this dataset is not affordable, almost a month and a half is needed, the evaluation was done using IsarPipeline only.

| number of seqs | isarPipeline runtime |
|---|---|
| 8817 | 170m 11s / 2h 50m |

**Table S1.** IsarPipeline runtime of running 8817 query batch, full TMSEG dataset without redundancy reduction, against a random sample of 68 million proteins from uniref90. As sequences are processed sequentially in PSI-BLAST, a multiplication by the average runtime of 11min would lead to about 1 month and half, roughly 540 times more than IsarPipeline runtime of 2h50m, to generate PSSM profiles.

The runtime needed to get the PSSM files via IsarPipeline is impressive (Tab. S1). We see that IsarPipeline needs only 170 minutes (Tab. S1) to generate the PSSMs while PSI-BLAST would take a month and a half. We observe a significant decrease in runtime by almost 540 folds.

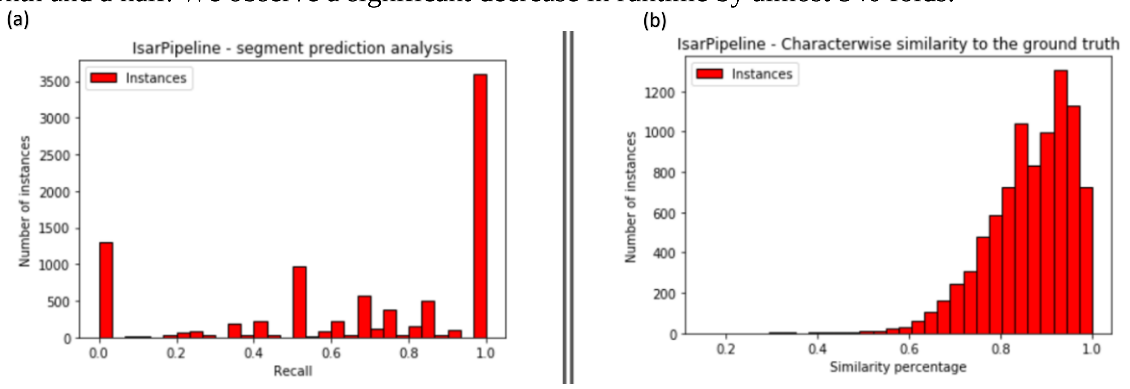


**Figure S6**. Analysis of PSSM-based predictions of the whole TMSEG dataset, without redundancy reduction. Histogram (a) represents the recall distribution, which is the percentage distribution of how many segments in a sequence ground truth are correctly predicted from an IsarPipeline PSSM. Histogram (b) represents the percentage distribution of how similar the residue predictions in a sequence are to the ground truth labels when using IsarPipeline PSSMs.

A full dataset analysis (Fig. S6) demonstrates that the distributions of both recall and characterwise percentage similarity are preserved. Additionally, we compare the overall recall and similarity means with the ones from previously evaluated sample. From a statistical stand point, IsarPipeline has a recall mean of **0.676** with a STD of **0.357**, which is a little bit higher than the sample one**.** Likewise, to the similarity mean, IsarPipeline has a similarity mean of **0.863** with a STD of **0.093**, which is again a bit higher than the sample statistic. For better statistical significance analysis, we convert all the previously reported standard deviations to standard errors (STE) using the following formula:

$$StdErr = \frac{StdDev}{\sqrt{Number\ of\ Samples}} \qquad \text{(Eq. I)}$$

| | **TMSEG Full Dataset Statistics (Fig. S6) – using IsarPipeline** | **TMSEG Sample Dataset Statistics (Fig. 3.a & S4.a) – using IsarPipeline** | **TMSEG Sample Dataset Statistics (Fig. 3.b & S4.b) – using PSI-BLAST** |
|---|---|---|---|
| Characterwise Analysis | 86.3% ± 0.01 | 83.0% ± 0.77 | 83.2% ± 0.73 |
| Segment-Based Analysis | 67.6% ± 0.38 | 60.7% ± 2.64 | 62.1% ± 2.55 |

**Table S2.** TMSEG statistical significance comparison of the segment-based analysis (via recall) and the characterwise analysis (via character percentage similarity). From a statistical significance comparison, IsarPipeline is as performant as PSI-BLAST on the sample dataset. The full dataset shows some kind of improvement, but this is just a feature of a larger dataset.

We observe in (Tab. S2) that with the full dataset we achieve higher performance using IsarPipeline compared to the sample analysis of TMSEG. However, this does not mean that IsarPipeline is outperforming PSI-BLAST since a similar analysis using PSI-BLAST is not conducted. The increase in performance can simply be argued to be a feature of the large database. Furthermore, the table (Tab. S2) depicts also that IsarPipeline performance was at least comparable, with interleaving performance, to PSI-BLAST results using the same sample test set.

### 4- IsarPipeline Evaluation on REPROF

#### 4.1 - Full Dataset Analysis

Similarly to TMSEG, we want to make sure that the sample we tested on is representative. We want to check whether the statistics computed previously still generalize for the whole REPROF dataset of 4692 protein sequences. Since running PSI-BLAST on this dataset is not affordable, almost three weeks will be needed, the evaluation was done using IsarPipeline only.

| number of seqs | isarPipeline runtime |
|---|---|
| 4692 | 73m 24s |

**Table S3.** IsarPipeline runtime required to run 4692 query batch, full REPROF dataset, against a random sample of 68 million proteins from uniref90. As sequences are processed sequentially in PSI-BLAST, a multiplication by the average runtime of 11min would lead to about 3 weeks, roughly 414 times more than IsarPipeline 73m, to generate PSSM profiles.

The runtime needed to get the PSSM files is again remarkable. IsarPipeline needs only 73m 24s (Tab. S3) to generate the PSSMs while PSI-BLAST would take about three weeks. A large decrease on runtime by almost 414 folds is achieved.

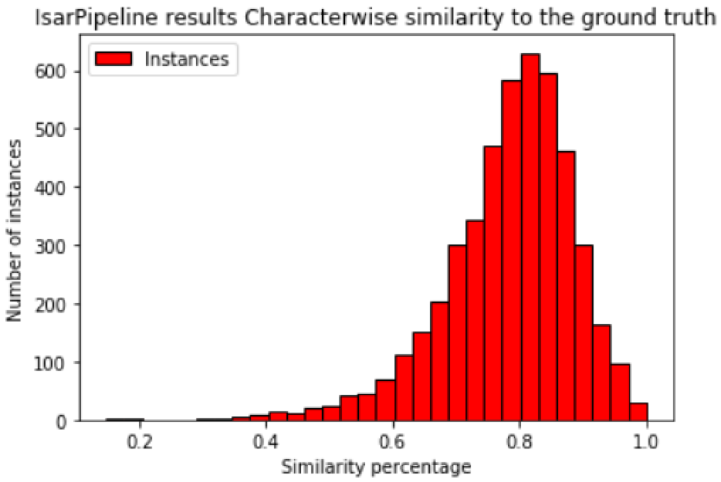


**Figure S7**. Analysis of PSSM-based predictions of the full REPROF dataset. The histogram represents the percentage distribution of how similar the secondary structure residue predictions in a sequence to the ground truth labels resulting from IsarPipeline PSSM files

The full dataset analysis (Fig. S7) demonstrates that the distribution of the characterwise percentage similarity is preserved. The histogram depicts a roughly sharp Gaussian distribution of the predictions around 0.8. From a statistical point of view, computing the overall similarity mean and comparing it to the previously evaluated sample, IsarPipeline has a similarity mean of **0.783** with a STD of **0.102**, which is a little bit higher than the sample statistic. For better statistical significance analysis, we convert all the previously reported standard deviations to standard errors using the formula in (Eq. I).

| | REPROF Full Dataset Statistics (Fig. S7) – using IsarPipeline | REPROF Sample Dataset Statistics (Fig. 5.a) – using IsarPipeline | REPROF Sample Dataset Statistics (Fig. 5.b) – using PSI-BLAST |
|---|---|---|---|
| Characterwise Analysis | 78.3% ± 0.15 | 78.1% ± 0.58 | 77.9% ± 0.58 |

**Table S4.** REPROF statistical significance comparison of the characterwise analysis (via character percentage similarity). Statistical significance of the three tests confirms that IsarPipeline performs generally the same as PSI-BLAST.

We clearly see (Tab. S4) that the performance did not change or been affected using IsarPipeline compared to PSI-BLAST only. We confirm, based on statistical significance, that IsarPipeline performance is comparable to PSI-BLAST results. Also, we can report through the full dataset analysis that our sample dataset is representative of the full one by preserving the statistics.
